# Aquaporin regulates cell rounding through vacuole formation during endothelial-to-hematopoietic transition

**DOI:** 10.1101/2022.09.03.506460

**Authors:** Yuki Sato, Mugiho Shigematsu, Maria Shibata-Kanno, Sho Maejima, Chie Tamura, Hirotaka Sakamoto

**Affiliations:** Graduate School of Medical Sciences, Kyushu University, Fukuoka, Japan; Ushimado Marine Institute, Graduate School of Natural Science and Technology, Okayama University, Setouchi, Okayama, Japan

**Keywords:** endothelial-to-hematopoietic transition, aquaporin, vacuole, cell rounding

## Abstract

Endothelial-to-hematopoietic transition (EHT) is crucial for hematopoietic stem cell (HSC) generation. During EHT, the morphology of hemogenic endothelial cells (HECs) changes from flat and adherent to spherical HSCs, which detach from the dorsal aorta. HECs attain a rounded shape in a mitosis-independent manner before cell adhesion termination, suggesting an atypical cell-rounding mechanism. However, the direct mechanisms underlying this change in cell morphology during EHT remain unclear. Here, we show that large vacuoles were transiently formed in HECs and that aquaporin-1 (AQP1) was localized in the vacuole and plasma membranes. Overexpression of AQP1 in non-HECs induced ectopic vacuole expansion, cell rounding, and subsequent cell detachment from the endothelium into the bloodstream, mimicking EHT. Loss of redundant AQP functions by CRISPR/Cas9 gene editing in HECs impeded the morphological EHT. Our findings provide the first evidence indicating that morphological segregation of HSCs from endothelial cells is regulated by water influx into vacuoles. These findings provide important insights for further exploration of the mechanisms underlying cell/tissue morphogenesis through water-adoptive cellular responses.

**SUMMARY STATEMENT:** Hemogenic endothelial cells transiently form large vacuoles during endothelial-to-hematopoietic transition. Aquaporin water channels regulate cell rounding and detachment of emerging hematopoietic stem cells through vacuole formation.

## INTRODUCTION

Hematopoietic stem cells (HSCs) arise directly from hemogenic endothelial cells (HECs) that are localized to the dorsal aorta floor during embryogenesis (Boisset et al., 2010; Kissa and Herbomel, 2010). Endothelial-to-hematopoietic transition (EHT) is a highly conserved transdifferentiation event leading to the generation of definitive HSCs in most vertebrate embryos (Dzierzak and Bigas, 2018; Klaus and Robin, 2017; North et al., 1999). Extrinsic signaling cues and transcriptional regulatory networks associated with HEC specification and HSC emergence have been closely studied (Dzierzak and Bigas, 2018; Wu and Hirschi, 2021); however, the mechanisms underlying the morphological transition from flat endothelial cells to round HSCs remain unclear. Runx1 is an essential transcription factor for EHT (Chen et al., 2009; Eliades et al., 2016; Howell et al., 2021; North et al., 1999). In fact, *Runx1*-deficient mouse embryos lack vacuole-like organelles with a spherical and ultralow-electron-density structure in their prospective HECs. These cells have further been noted to exhibit irregular cell flattening (North et al., 1999). This suggests the involvement of vacuoles in the process of cell rounding; however, the mechanisms underlying their generation as well as their biological roles in HECs remain unclear.

In plants, vacuoles play a pivotal role in the regulation of cell morphology and size in response to osmotic pressure. Aquaporin (AQP) family proteins mediate water transport in both cell and vacuole membranes (Maurel et al., 2015). In animal models, water influx through AQP1 promotes migration of vascular endothelial cells, cancer cells, bone marrow mesenchymal cells, and neural crest cells (De Ieso and Yool, 2018; Huebert et al., 2010; Kao et al., 2017; McLennan et al., 2020; Meng et al., 2014; Saadoun et al., 2005). Meanwhile, AQP1 expression and channel function are regulated by osmotic and hydrostatic pressure, fluid shear stress, and hypoxia (Huo et al., 2021; Kao et al., 2017; Morishita et al., 2019; Nguyen et al., 2015; Verkman, 2002) Given that HECs in the dorsal aorta directly contact circulating blood, they are more likely to directly take up water in response to these mechanophysiological stimuli.

During EHT in zebrafish embryos, HECs begin rounding toward the basal side of the dorsal aorta endothelium; subsequently, HSCs are extruded into the sub-aortic space. In this process, apical membrane invagination and subsequent cavity formation through circumferential actomyosin contraction are accompanied by cell rounding (Lancino et al., 2018). In contrast, within amniote embryos, HECs begin rounding toward the apical side, and HSCs are released directly into the blood stream (Boisset et al., 2010). Hence, the cellular mechanisms that control morphological changes during EHT in amniote embryos might be different from those activated in fish. Avian embryos serve as an appropriate model system for hematopoietic research because they readily undergo various embryo manipulations, including tissue transplantation, exogenous gene transduction, and live imaging (Jaffredo et al., 2010; Kulesa et al., 2013; Asai et al., 2020). In the current study, we sought to better understand the regulatory mechanisms underlying morphological transition from HECs to HSCs in amniotes. To this end, we characterized vacuoles in HECs using chick and quail embryo models, analyzed AQP1 localization, and proposed a simple model of endothelial cell rounding by water permeation.

## RESULTS

### Vacuoles are specifically formed during EHT

In this study, we selected chick or quail embryos depending on the availability of genetic resources, markers, and live imaging (Motono et al., 2010; Pardanaud et al., 1987; Sato et al., 2010). Based on a report that untagged fluorescent proteins are excluded from vacuoles (Garcia et al., 2017), we utilized transgenic (tg) chick embryos, tg(pLSiΔAeGFP) (Motono et al., 2010), which express eGFP ubiquitously, to quantitatively characterize the vacuoles. Runx1-positive HECs were found in the dorsal aortic floor from embryonic day (E) 2.5 to 4 (Fig. 1A–C, E). During this period, eGFP-negative regions were observed in the HECs. Concurrent with EHT completion at E5, endothelial cells containing eGFP-negative regions were rarely observed (Fig. 1D–F). Therefore, vacuoles in the dorsal aortic floor were identified as transient organelles that may be found in HECs. Use of Tg(pLSiΔAeGFP) embryos allowed the measurement of vacuole sizes. HECs in the floor exhibited various vacuole sizes, which were significantly larger, on average, than those in non-HECs in the roof (Fig. 1G–I). Commonly observed small vesicles in both the floor and roof were endosomes yielded by the intrinsic endocytic activity of endothelial cells (Simionescu *et al*., 2002).

**Figure 1.**
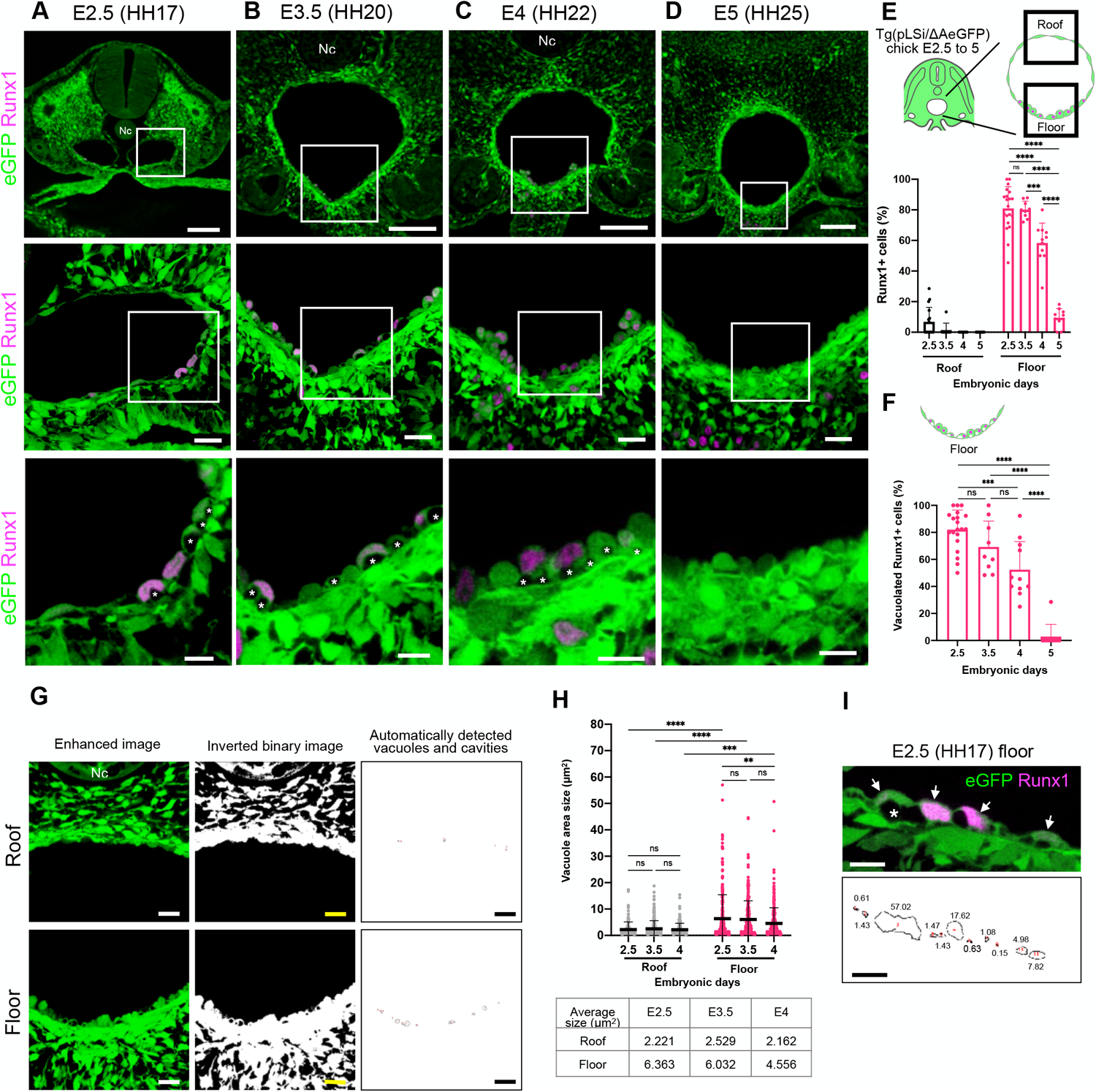
Vacuoles are transiently observed in the dorsal aorta during EHT. Optical cross-sections of tg(pLSiΔAeGFP) chick embryos at E2.5 (A), 3.5 (B), 4 (C), and 5 (D). The magnified areas are indicated by squares. HECs were detected using anti-Runx1 immunostaining. Vacuoles are identified with asterisks. Runx1^+^ cells underneath the dorsal aorta were hematopoietic cells stored in para-aortic foci (B–D) (Fellah et al., 2013). (E) Percentage of Runx1^+^ cells in the roof and floor endothelial cells of the dorsal aorta. (F) Percentage of vacuolated Runx1^+^ cells in each stage. (G) Inverted binary images of eGFP were used for the automatic segmentation and measurement of vacuole sizes. (H) Cross-sectional area sizes of the vacuoles in the dorsal aortic roofs and floors at E2.5, 3.5, 4, and 5. (I) Representative images of the eGFP-negative areas observed in the floor on E2.5. Outlines of the inverted images were drawn using ImageJ software (lower panel). Each value indicates eGFP-negative area sizes (μm^2^). Runx1^+^ cells are indicated by the arrows. Extremely large eGFP-negative areas of over 30 μm^2^ are cavities spanning multiple cells (asterisk); 20 slices, n = 3, at E2.5; 9 slices, n = 3 at E3.5; 11 slices, n = 3 at E4; and 10 slices, n = 3 at E5 were quantified in (E and F). 6 slices, n = 3 at E2.5; 9 slices, n = 3 at E3.5; and 9 slices, n = 3 at E4 were quantified in (H). Error bars indicate standard deviation (SD), ns: not significant, *** P < 0.0005, **** P < 0.0001 by one-way ANOVA. Nc: notochord. Scale bars: 100 μm in the upper panel, 20 μm in the middle panel, and 10 μm in the lower panel in (A). 20 μm in (G) and 10 μm in (I).

### AQP1 is localized in the plasma and vacuole membranes

We found that AQP1 was localized to both vacuoles and plasma membranes in the endothelial cells during EHT (E2.5 to 4, Figs 2A–D, S1A–B); its expression decreased following EHT completion (E5, Fig. S1C–F). AQP1 expression was observed in endothelial cells of the dorsal aorta, regardless of dorsoventral position (Fig. 2B). Accordingly, AQP1 expression was not restricted to Runx1-positive HECs (Fig. 2E). This suggests that AQP1 expression is regulated independently of Runx1; however, AQP1 downregulation by E5 occurred simultaneously with the disappearance of Runx1-expressing HECs (Fig. S1F).

**Figure 2.**
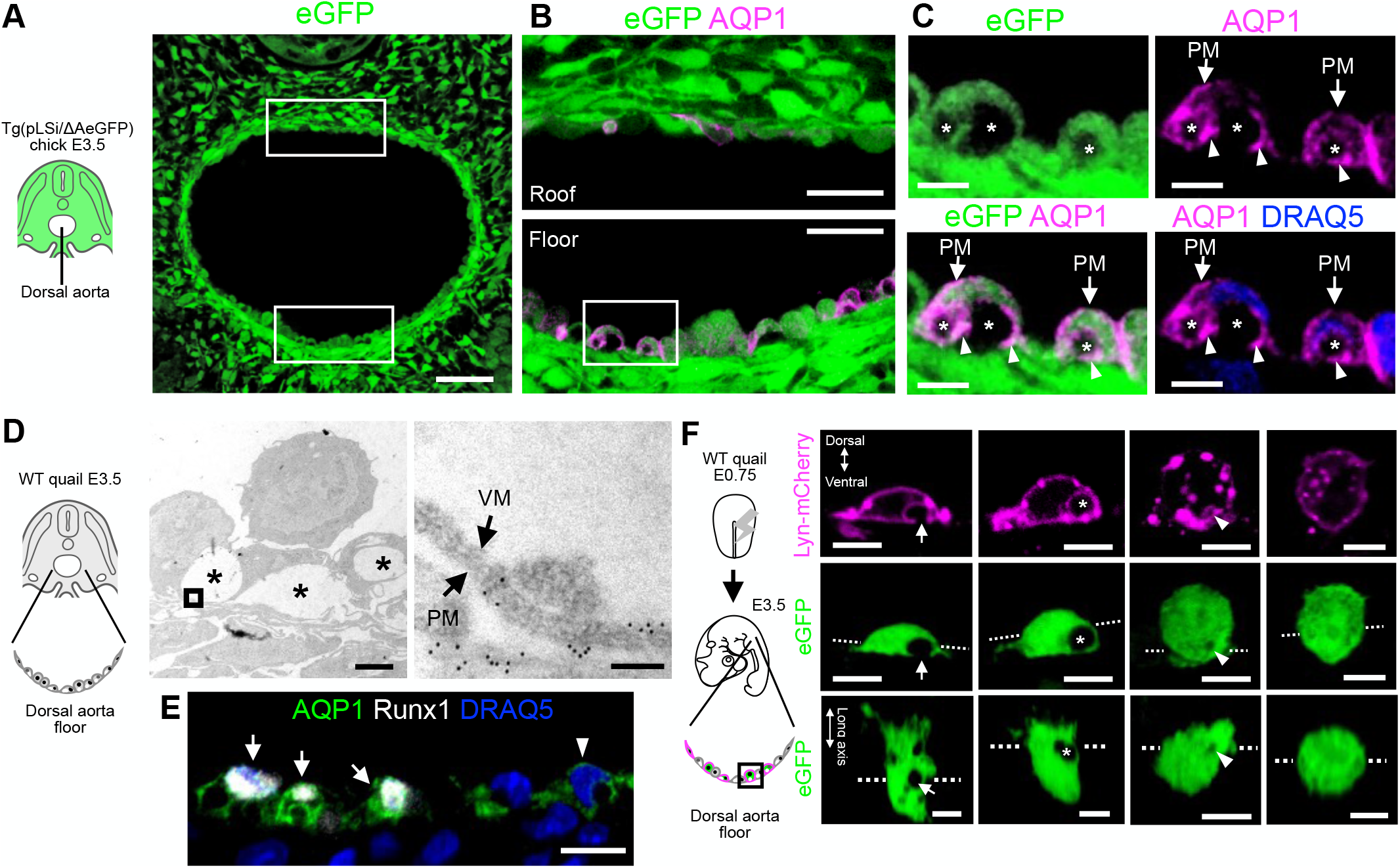
AQP1 is localized in the plasma and vacuole membranes of endothelial cells in the ventral side of the dorsal aorta. (A–C) Optical cross-sections of a tg(pLSiΔAeGFP) embryo at E3.5. (B) Enlarged view of the rectangular areas in (A). (C) Enlarged view of rectangular areas in (B). Endothelial cells in the dorsal aortic floor show eGFP-negative areas (asterisks). AQP1 is localized in the plasma membrane (PM) and boundaries of the eGFP-positive and -negative areas (arrowheads). (D) Immunoelectron microscopy of wild-type (WT) quail embryos at E3.5. Endothelial cells in the dorsal aortic floor contain large vacuoles (asterisks). An enlarged view of the square area is shown in the right panel. AQP1 (6-nm gold particles) was localized in the plasma and vacuole membrane (VM). (E) Optical cross-sections of the dorsal aortic floor in a WT quail embryo at E3.5. AQP1 is localized in the plasma and vacuole membranes in Runx1-positive (arrows) and –negative (arrowheads) cells. (F) Expression vectors for eGFP and Lyn-mCherry were co-electroporated into progenitor cells on the primitive streak at E0.75 (HH4). Optical cross-sections of the dorsal aortic floor (upper two panels) in electroporated quail embryos at E3.5. In a hemispherical cell, an eGFP-negative cavity was observed on the basal side (arrows). In another hemispherical cell, eGFP-negative vacuoles (asterisks) delineated by a Lyn-mCherry-positive membrane were completely separated from the PM. A small vacuole was observed in the rounded cell (arrowheads). Lower: Optically reconstructed horizontal sections. Dashed lines indicate optical slice positions. Scale bars: 50 μm in (A), 20 μm in (B), 5 μm in (C, E), 2 μm in (D, left), 100 nm in (D, right), 5 μm in (E), and 10 μm in (F).

To visualize individual cell morphologies, dorsal aortic endothelial cells were labeled with Lyn-mCherry and eGFP by electroporation (Fig. 2F). An additional eGFP-negative structure was observed that was not fully detached from the plasma membrane. We identified this structural cavity as being distinct from the vacuole. Meanwhile, vacuoles were observed in the hemispherical and nearly spherical cells but were not detected in fully rounded cells (Fig. 2F). We predicted that the cavity was a precursor of the vacuole, as most of the vacuoles were found in hemispherical cells. Based on these results, we hypothesize that AQP1 plays a role in the cell-rounding process of HECs by mediating water permeation into cavities and vacuoles.

### AQP1 promotes ectopic vacuole formation and cell rounding

The water permeability of AQP family proteins has been studied using cell swelling analyses of *Xenopus* oocytes by overexpressing AQP proteins in various species (Preston *et al*., 1992; Beitz *et al*., 2009). Hence, we examined the effects of overexpressing AQP1 in non-HECs. To this end, the AQP1-mRFP expression vector was electroporated into an aortic roof-specific cell lineage (Pardanaud et al., 1996). Vacuoles were detected in the co-electroporated eGFP-negative areas. AQP1-mRFP-overexpressing cells had expanded large vacuoles ectopically on the roof (Fig. 3A and B). Moreover, quantification of the eGFP-negative area sizes indicated that AQP1 overexpression induced a marked increase in vacuole size (Fig. 3C). Meanwhile, Runx1 expression was not detected in these cells, suggesting that vacuole expansion was Runx1-independent.

**Figure 3.**
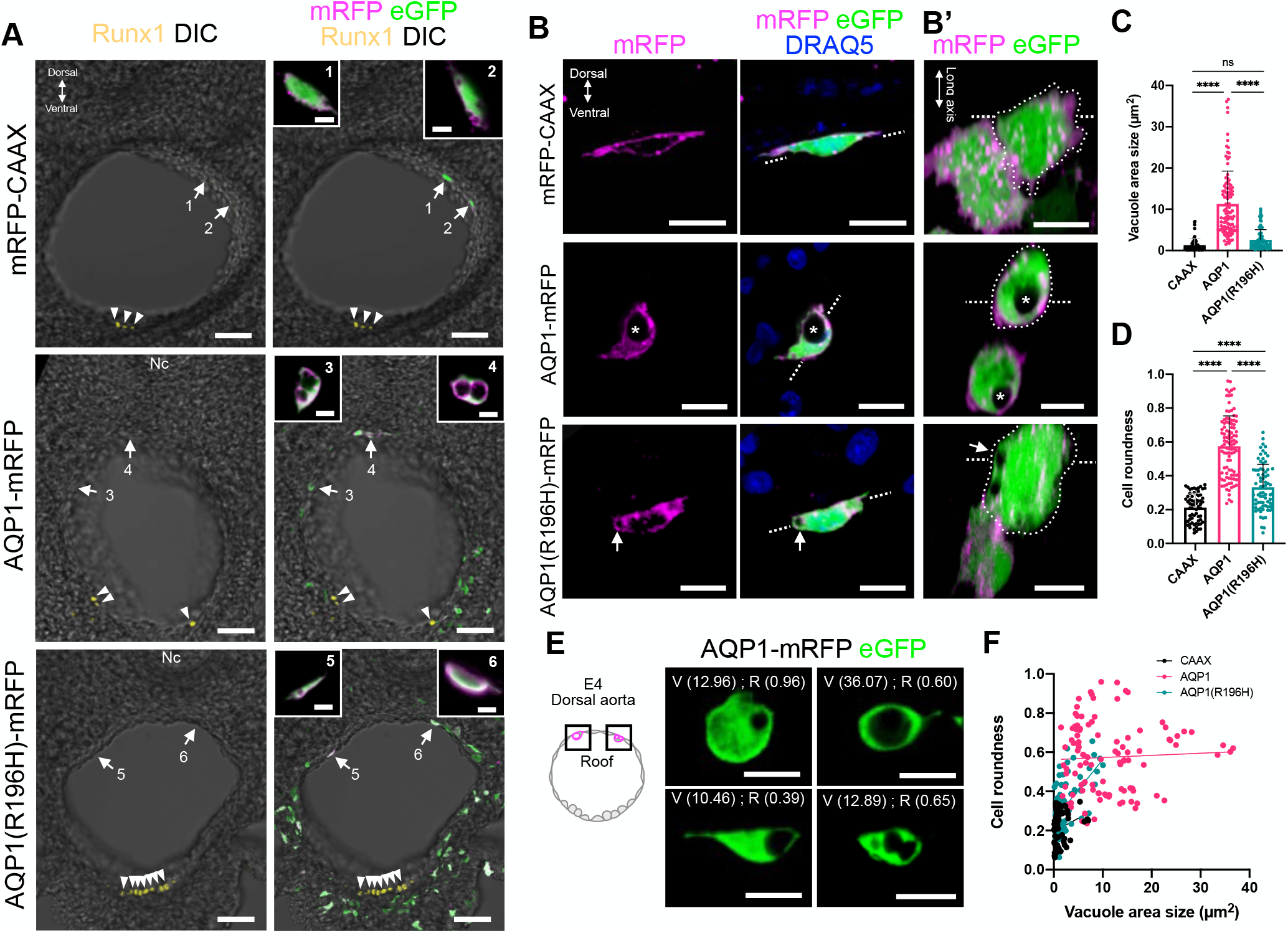
AQP1 overexpression leads to ectopic vacuole expansion and cell rounding. (A) Optical cross-sections of mCherry-CAAX (control)-, AQP1-mRFP-, and AQP1(R196H)-mRFP-overexpressing quail embryos at E4. eGFP was coexpressed for vacuole measurements. Runx1-positive HECs are indicated by arrowheads. AQP1-mRFP-overexpressing endothelial cells ectopically formed large vacuoles without Runx1 expression in the roof of the dorsal aorta (cells 3 and 4 in the middle panels). Large vacuoles were not observed in AQP1(R196H)-mRFP-overexpressing endothelial cells (cells 5 and 6 in the lower panels). (B) Representative optical cross- (B) and horizontal (B’) sections of mCherry-CAAX-, AQP1-mRFP-, and AQP1 (R196H)-mRFP-overexpressing cells in the roof. Thick dashed lines indicate optical slice positions. A large vacuole was formed ectopically in AQP1-mRFP-overexpressing cells (asterisks in the middle panels). In AQP1(R196H)-mRFP-overexpressing cells, vacuoles were smaller than those in AQP1-mRFP-overexpressing cells (arrows in the lower panels). (C) Sizes of the cross-sectional areas of vacuoles in the roof. AQP1-mRFP-overexpressing cells formed significantly larger vacuoles. (D) Roundness of vacuolated cell cross-sections. AQP1-mRFP-overexpressing cells were rounded compared with mRFP-CAAX- and AQP1(R196H)-mRFP-overexpressing cells. (E) Representative images of co-electroporated eGFP in AQP1-mRFP-overexpressing cells in the roof. Values in brackets are the vacuole size (V) and cell roundness (R). (F) Scatter plot of vacuole size and cell roundness shown in (C and D). AQP1-mRFP-overexpressing cells were characterized by large vacuoles and increased roundness. The correlation coefficients were r = 0.2466 (p = 0.0459) for mRFP-CAAX, r = 0.04878 (p = 0.6229) for AQP1-mRFP, and r = 0.5672 (p<0.0001) for AQP1(R196H)-mRFP; 66 cells, n = 9 of mRFP-CAAX; 104 cells. n = 9 of AQP1-mRFP; 75 cells, n = 9 of AQP1(R196H)-mRFP-overexpressing embryos were quantified. Error bars indicate SD. ns: not significant; ****p<0.0001 using one-way ANOVA. Nc: notochord. Scale bars: 50 μm in (A), 10 μm in (B), and 5 μm in (F).

To determine whether the selective water transport ability of AQP1 is responsible for vacuole expansion, an AQP1 mutant for the aromatic/arginine constriction region (ar/R), AQP1(R196H)-mRFP, was overexpressed (Beitz et al., 2009; Khan *et al*., 2013). AQP1(R196H)- mRFP-expressing cells formed small vacuoles (Figs 3A and B), comparable in size to those of the control lipid membrane reporter, mRFP-CAAX-overexpressing cells (Fig. 3C), indicating that water transport into the vacuoles facilitates their expansion in endothelial cells. Quantification analysis further revealed that AQP1-mRFP-overexpressing cells were significantly rounded compared to mRFP-CAAX- and AQP1(R196H)-mRFP-overexpressing cells (Fig. 3D). These results indicate a simple mechanism by which water influx into the vacuole promotes cell rounding.

However, large vacuoles were not observed in the fully spherical cells (roundness > 0.8). In contrast, hemispherical cells (roundness 0.5–0.7) possessed large vacuoles (Fig. 3E). Indeed, the scatter plot indicated no significant linear relationship between vacuole size and cell roundness in AQP1-overexpressing cells (Fig. 3F). Thus, the vacuole expansion induced by excess AQP1 contributes to the flat-to-hemispherical transition phase. Similar to vacuoles in normal HECs, these artificially expanded vacuoles later regressed through some intrinsic degradation mechanism during the completion of ectopic cell rounding.

### Ectopically rounded cells enter the circulation

To confirm whether ectopically rounded cells detach from the vascular endothelium, we performed time-lapse imaging of AQP1-overexpressing quail embryos. In most amniote embryos, during EHT stages, the dorsal aorta is located deep in the trunk, thus impeding live *in vivo* imaging analysis of EHT. Therefore, an optically accessible non-hemogenic vasculature of proximal vitelline vessels at E2 was selected for analysis (Fig. 4A). In AQP1-2A-eYFP-overexpressing embryos, eYFP-positive endothelial cells in the vitelline vessels became rounded within a few hours (Fig. 4B). The ectopically rounded cells began to flow downstream and subsequently disappeared from the image field (Movie 1). Hence, non-HEC rounding caused by excess AQP1 expression was accompanied by ectopic cell detachment from the endothelium. Moreover, in the magnified views, the eGFP-labeled control cells displayed an elongated shape with continuous migratory activity and no detectable vacuoles (Fig. 4C and Movie 2). In contrast, the AQP1-2A-eYFP-overexpressing cells displayed vacuoles (Fig. 4C and Movie 2) that became undetectable as cell rounding progressed. The rounded cells eventually disappeared from the vitelline vessels. At this stage, the number of apoptotic cells had not increased in the AQP1-2A-H2B-mCherry-overexpressing regions (Fig. S2A and B) and no abnormal nuclear morphology or DNA punctum was observed in the largely vacuolated cells (Fig. S2C). Therefore, detachment of the AQP1-overexpressing cells did not associate with apoptotic cell extrusion, and artificially formed vacuoles were not cytotoxic. Such ectopic vacuole formation induced by AQP1 overexpression was not observed in the surrounding mesenchymal cells outside the vitelline vessels (Fig. S2D). These AQP1-overexpressing cells continued circulating and were observed in the blood stream at a later stage (Movie 3). Hence, vacuole expansion induced by excess AQP1 promotes endothelial cell rounding and subsequent detachment from the vascular endothelium.

**Figure 4.**
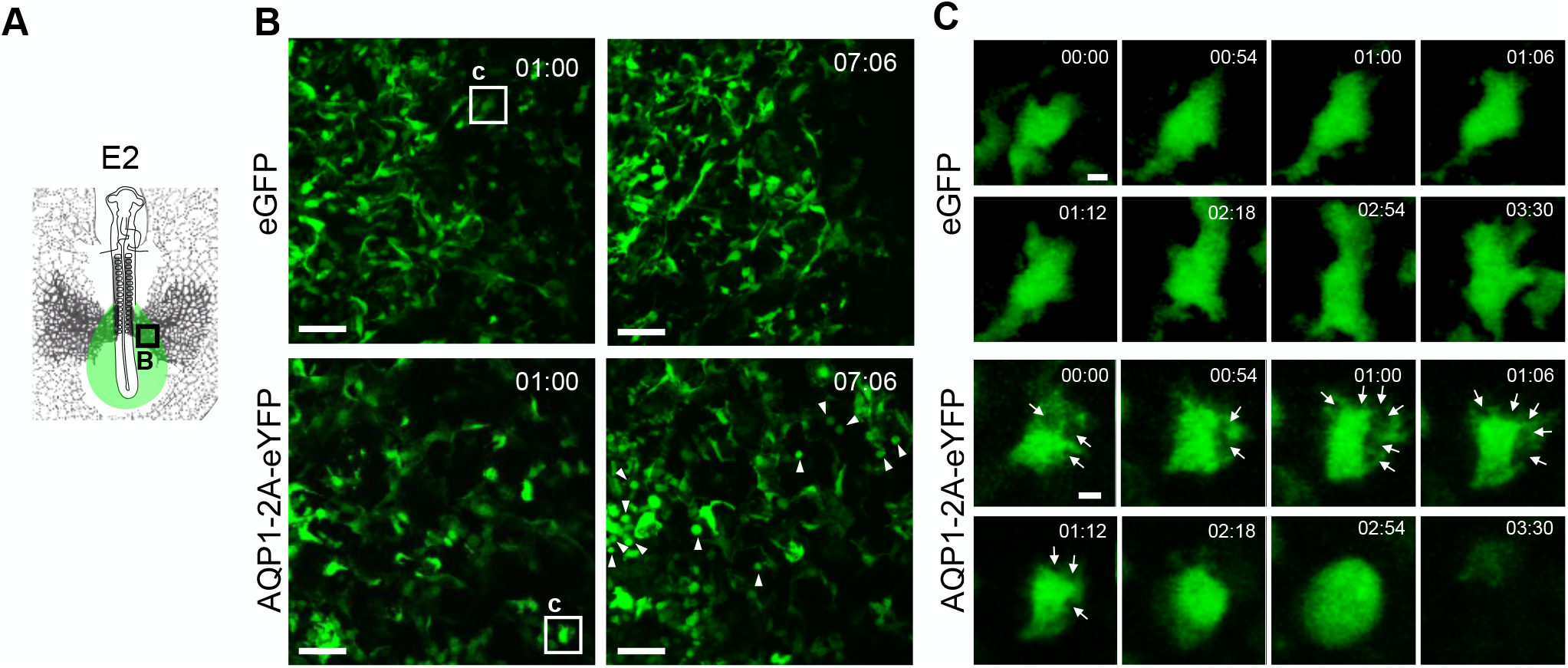
AQP1-mediated ectopic vacuole formation is accompanied by cell detachment. (A) Live imaging analysis of ectopic EHT in quail embryos. The electroporated region is shown in green. Squares indicate the imaged areas shown in (B). (B) Time-lapse observation of vitelline vessels in eGFP (control)- and AQP1-2A-eYFP-overexpressing embryos. AQP1-2A-eYFP-overexpressing endothelial cells became rounded and detached from vitelline vessels (arrowheads in lower panels, see Movie 1). Squares in the left panels indicate enlarged cells in (C). (C) Control cells did not form detectable vacuoles (upper panels). In AQP1-2A-eYFP-overexpressing cells, vacuoles (arrows) formed concurrently with cell rounding. Eventually, AQP1-2A-eYFP-overexpressing cells were detached (see Movie 2). Optical horizontal sections (B and C), Scale bars: 50 μm in (B), 5 μm in (C).

### Hemogenic endothelial cells form vacuoles *in vitro*

*In vitro* reconstitution of EHT is possible in avian embryos (Yvernogeau et al., 2016). To investigate whether vacuoles and cavities are formed *in vitro*, we cultured the quail embryo-derived presomitic mesoderm (PSM, Fig. 5A). Within a few days, the QH1-positive endothelial cells were rounded and expressed AQP1 (Fig. 5B). Culture of the tg(pLSiΔAeGFP) embryo-derived PSM revealed that Runx1-positive HECs formed cavities and vacuoles (Fig. 5C and D). Meanwhile, AQP1 was localized in the plasma and vacuole/cavity membranes, similar to that observed in *in vivo* HECs (Fig. 5E). These results suggest that vacuoles are formed concurrently with EHT under *in vitro* conditions.

**Figure 5.**
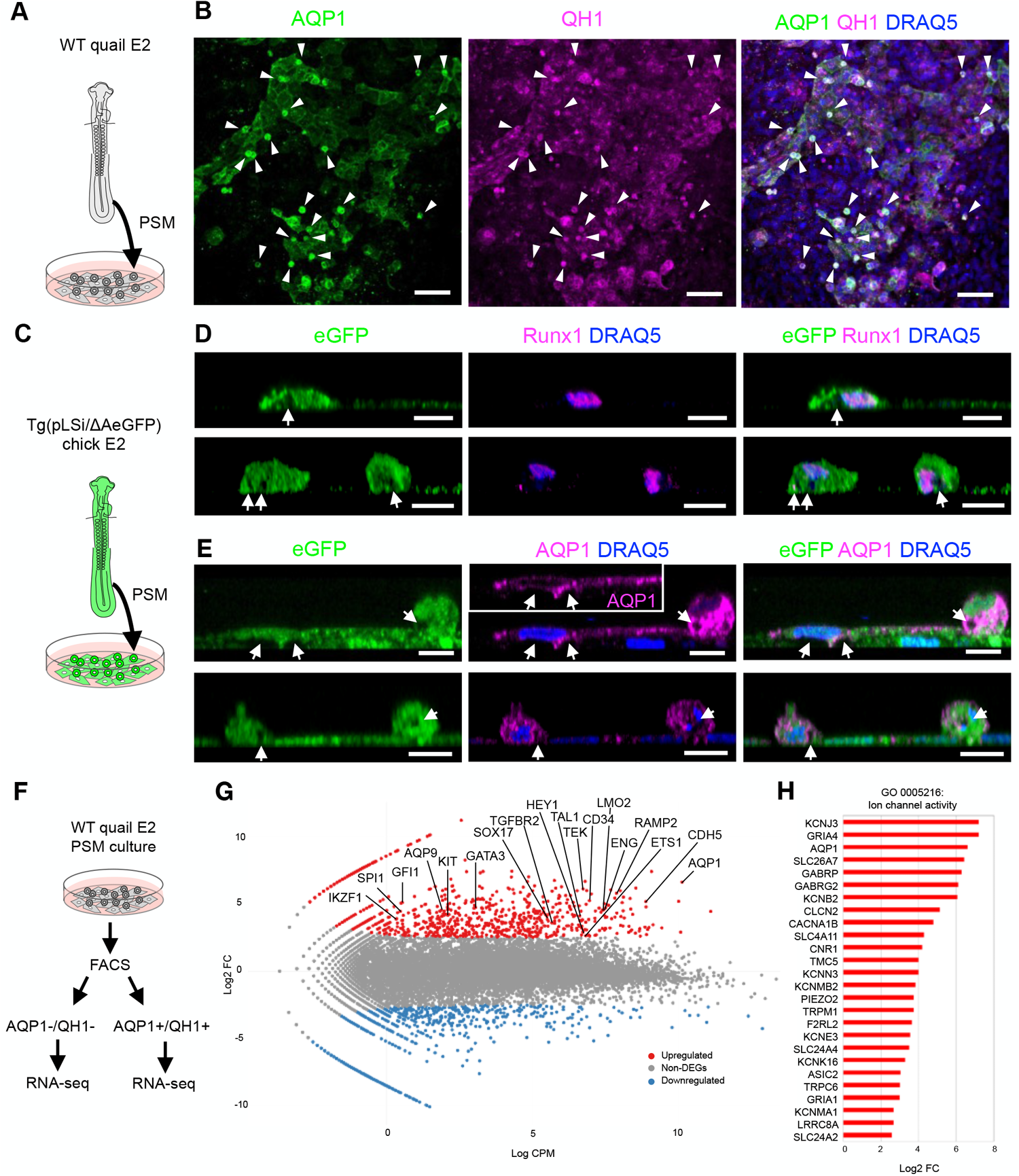
*in vitro* EHT recapitulates AQP1 localization in vacuoles/cavities. (A) EHT in cultures of the presomitic mesoderm (PSM) isolated from quail embryos (Yvernogeau et al., 2016). (B) Z-stacked frontal images of cultured tissues at 72 h. AQP1 is expressed in QH1-positive endothelial cells (Pardanaud et al., 1987). Rounded cells are indicated by arrowheads. (C) Vacuole/cavity formation in tg(pLSiΔAeGFP)-derived tissue. (D and E), Reconstructed optical cross-sections of tg(pLSiΔAeGFP)-derived tissues at 72 h. eGFP-negative vacuoles and cavities (arrows) formed in Runx1-positive hemogenic endothelial cells (D). AQP1 localization in the plasma and vacuole/cavity membranes (E). (F) RNA-seq for AQP1^+^/QH1+ and AQP1^−^/QH1^−^ cells at 72 h. (G) MA plot of differentially expressed genes (DEGs). X and Y axes are Log CPM (count per million) and Log2 FC (fold change), respectively. Log2 FC > 1, P < 0.05 and Log2 FC < −1, P < 0.05 are indicated by red and blue, respectively. Embryonic hematopoiesis-related genes and AQPs are identified. (H) Upregulated genes associated with ion channel activity in the AQP1^+^/QH1^+^ population. Scale bars: 50 μm (B) and 10 μm (D and E).

To characterize AQP1-expressing endothelial cells, AQP1/QH1 double-positive cells and AQP1/QH1 double-negative cells were sorted from cultured tissues and subjected to RNA-seq analysis (Fig. 5F). Differentially expressed gene (DEG) analysis of these two different cell populations revealed that genes involved in the specification of HECs and HSCs were upregulated in AQP1/QH1 double-positive cells (Fig. 5G). Based on the enriched gene ontology (GO) terms, the AQP1/QH1 double-positive cell population was characterized as vascular endothelial cells that potentially differentiated into hematopoietic cell lineages (Fig. S3). In this population, various ion channels and transporters were transcriptionally upregulated (Fig. 5H). Among them, proteins encoded by *TRPM1, TRPC6*, and *PIEZO2* mediate mechanical stimulus (GO: 0009612, Negri et al., 2020), and those encoded by *SLC4A11, KCNMA1*, and *LRRC8A* mediate the response to osmotic stress (GO: 0006970). These molecules are expected to be involved in the regulation of local osmolarity in HECs (discussed later).

### AQP channels are redundantly required for HEC rounding

Abnormalities in EHT have not been reported in AQP1-deficient mouse embryos (Hua et al., 2019). Therefore, to assess the requirement of AQP1 for EHT in quail embryos, we used CRISPR/Cas9 genome editing for *AQP1* knockout. AQP1-deficient endothelial cells—confirmed by loss of AQP1 protein expression—showed large vacuoles and rounded cell shapes (Fig. S4). We hypothesized that AQP1 is not solely involved in EHT, as AQP5, AQP8, and AQP9 were also redundantly expressed in the endothelium of the dorsal aorta (Fig. S5A). Since AQP9 was upregulated in AQP1-expressing endothelial cells *in vitro* (Fig. 5G), double knockout analyses were performed; however, co-electroporation of AQP1 and AQP9 guide RNA (gRNA) with Cas9-2A-mRFP in HECs did not cause obvious deficiency in vacuole formation or cell rounding. Similarly, double-knockouts of *AQP1* and *AQP5* as well as *AQP1* and *AQP8* did not induce any morphological abnormalities (Fig. S5B and C).

CRISPR/Cas9 genome editing by electroporation cannot attain 100% efficiency (Williams et al., 2018). We, therefore, aimed to achieve the stochastic reduction of functional AQPs in HECs by introducing multiple gRNAs for *AQP1, AQP5, AQP8*, and *AQP9* with Cas9-2A-mRFP. In electroporated embryos, multiple *AQP*-knockout HECs (mRFP^+^/Runx1^+^) exhibited a flat morphology (Fig. 6A and B). In these embryos, the percentage of vacuolated HECs significantly decreased (Fig. 6C). Among the vacuolated cell population, notable reductions in vacuole area size and cell roundness were observed (Fig. 6D–F). Accordingly, non-vacuolated cells exhibited lower cell roundness (Fig. 6D and F). These results indicate that redundant AQP activities are required for vacuole expansion and cell rounding.

**Figure 6.**
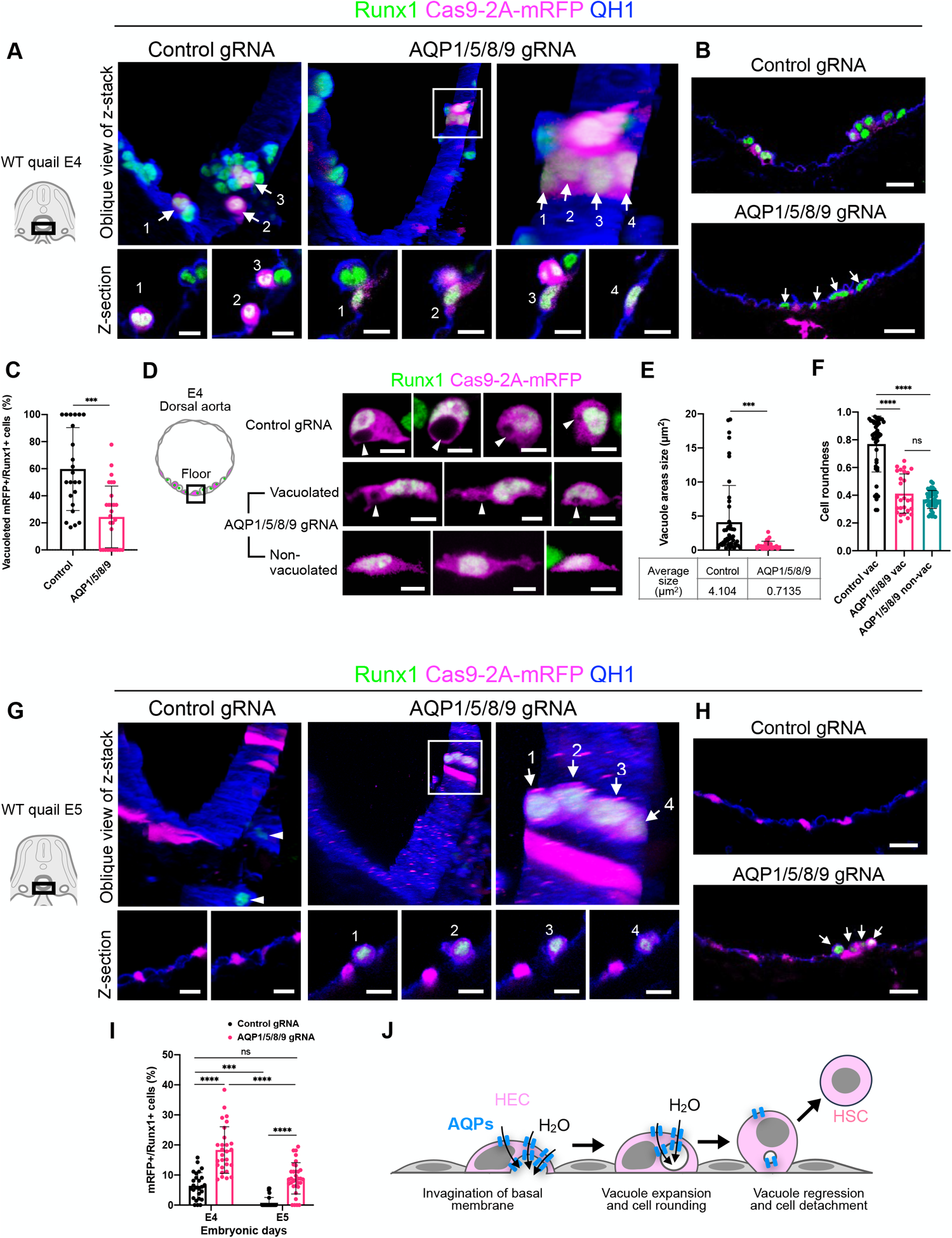
Redundant AQP function is required for hemogenic endothelial cell rounding and detachment. (A) Oblique views of z-stacked images of the dorsal aortic floor at E4 (upper panels). Electroporated cell bodies and internal vacuoles were identified by mRFP. Endothelial cell membrane is detected by QH1. Arrows in the left panel: mRFP^+^/Runx1^+^ cells in control gRNA-electroporated embryos. Arrows in the right panel: mRFP^+^/Runx1^+^ cells in AQP1/5/8/9 gRNA-electroporated embryos. Lower panels: optical cross-sections of each mRFP^+^/Runx1^+^ cell. (B) Optical cross-sections of the aortic floor at E4. Upper: mRFP^+^/Runx1^+^ cells in control embryos. Lower: mRFP^+^/Runx1^+^ cells in AQP1/5/8/9 gRNA-electroporated embryos. Arrows: mRFP^+^/Runx1^+^ flat cells. (C) Percentage of vacuolated mRFP^+^/Runx1^+^ cells among the total Runx1^+^ cells in the control (23 slices, n = 13) and AQP1/5/8/9 gRNA-electroporated embryos (28 slices, n = 14) at E4. (D) Enlarged views of representative electroporated HECs (mRFP^+^/Runx1^+^). Vacuoles are indicated by arrowheads. (E) Cross-sectional area sizes of the vacuoles in control (46 cells, n = 13) and AQP1/5/8/9 gRNA-electroporated embryos (27 cells, n = 14) at E4. (F) Cell roundness of mRFP^+^/Runx1^+^ cells in control (46 cells, n = 13) and AQP1/5/8/9 gRNA-electroporated embryos at E4 (27 vacuolated and 46 non-vacuolated cells, n = 14). (G) Oblique views of z-stacked images of the dorsal aortic floor at E5 (upper panels). Left: mRFP^+^/Runx1^+^ cells in control embryos. Arrowheads: Runx1^+^ cells. Arrows: mRFP^+^/Runx1^+^ cells in AQP1/5/8/9 gRNA-electroporated embryos. Lower: optical cross-sections of each mRFP^+^/Runx1^+^ cell. (H) Optical cross-sections at E5. Arrows: mRFP^+^/Runx1^+^ cells in AQP1/5/8/9 gRNA-electroporated embryo. (I) Percentages of mRFP^+^/Runx1^+^ cells among eYFP+ endothelial cells in the dorsal aortic floors of control gRNA (26 slices, n = 5 at E4; 30 slices, n = 5 at E5) and AQP1/5/8/9 gRNA (28 slices, n = 6 at E4; 33 slices, n = 9 at E5) electroporated tg(tie1:H2B-eYFP) embryos (refer to Fig. S6). (J) Model depicting HEC rounding. Error bars indicate SD. ns: not significant, *** P < 0.001, **** P < 0.0001 by unpaired t-test (C and E) and one-way ANOVA (F and I). Scale bars: 10 μm (lower panels in A and G), 20 μm (B and H) and 5 μm (D).

As we noted previously (Fig. 1), EHT was completed by E5, and Runx1-positive HECs disappeared from the dorsal aorta. Consistently, in control gRNA-electroporated embryos, Runx1-positive HECs were not found in the dorsal aorta at E5 (Fig. 6G and H). In contrast, multiple *AQP*-knockout HECs were detected in the floor at this stage. Tg(tie1:H2B-eYFP) quail embryos fluorescently label endothelial cells and HECs, allowing the accurate segmentation of individual cell nuclei and subsequent automatic cell counting (Sato et al., 2010). Quantification of the mRFP^+^/Runx1^+^ cells in tg(tie1:H2B-eYFP) embryos revealed that multiple *AQP*-knockout HECs were selectively retained in the floor of dorsal aorta at E5 (Figs 6I and S6), indicating that the detachment of HECs from the endothelium was suppressed by multiple *AQP* knockouts. We, therefore, concluded that cell rounding and detachment during EHT are regulated by AQP-mediated water permeation into vacuoles (Fig. 6J). As it has been suggested that AQP1 transitions from a high-water-permeability state to a closed state in response to an increase in membrane tension without osmotic changes (Ozu et al., 2013), AQPs in rounded HECs may automatically stop water transport and prevent plasma membrane rupture.

## DISCUSSION

Our results suggest that vacuole expansion, cell rounding, and detachment of endothelial cells by AQP are Runx1-independent processes (Figs 3 and 6). A previous genome-wide search for Runx1 binding sequences in cultured mouse HECs implied that AQP genes are not involved in direct transcriptional targets of Runx1 (Lie-A-Ling et al., 2014). However, Runx1 is genetically required for vacuole formation (North et al., 1999), and its expression is correlated with the progression of cell rounding in mouse embryos (Bos et al., 2015). Runx1 also regulates the expression of genes involved in cytoskeletal reorganization (Lie-A-Ling et al., 2014). In our study, HECs with multiple *AQP-*knockout exhibited flattened shapes with Runx1 expression detected at E4; however, these cells were hemispherically raised from the endothelium at E5 (Fig. 6). This implies that morphological changes in HECs are not solely dependent on AQP function. We, therefore, hypothesized that during normal development, AQP-mediated water permeation synergistically promotes EHT with downstream targets of Runx1.

Osmotic and hydrostatic pressure are involved in the opening/closing of the AQP channel (Nguyen et al., 2015; Verkman, 2002). Nonetheless, the mechanophysiological conditions and molecules involved in the regulation of AQP functions in HECs remain to be elucidated. We demonstrated that excess AQP1 expression induces ectopic EHT-like cellular responses without experimental changes in osmotic and hydrostatic pressures (Figs 3 and 4). Although single *AQP1-* knockout HECs retained their vacuole-formation and cell-rounding abilities, multiple knockouts of redundantly expressing *AQP1, AQP5, AQP8*, and *AQP9* genes in HECs led to failure of morphological EHT (Figs 6 and S4). These results indicate that the endothelial cell-rounding process is AQP dose-dependent. Moreover, in the AQP1-expressing endothelial cells *in vitro*, a considerable number of ion channels and transporters, which may contribute to osmoregulation and/or mechano-transduction, were upregulated (Fig. 5). We, therefore, searched for deletion phenotypes of these genes; however, none of them were found to influence embryonic hematopoiesis. Generally, ion channels and transporters belong to a large family of genes with overlapping functions. Hence, the mechanophysiological activation mechanisms of AQPs require further investigation in light of the functional redundancy of these solute channels/transporters, which may have been acquired to tightly regulate EHT.

AQP1 localization suggests the possibility that water accumulation begins in the basal space of HECs, which triggers the invagination of the basal membrane followed by internalization of cavities into the vacuoles. Based on our observation, we speculate that water accumulation on the basal side mechanically bends the plasma membrane, thereby facilitating membrane invagination and subsequent large vacuole formation in HECs. However, vacuoles eventually exhibit reduction in their average size and disappear from fully rounded HECs (Figs 1 and 2). Consistently, time-lapse imaging analysis revealed that the ectopically induced vacuoles quickly became undetectable prior to cell detachment (Fig. 4). Knowledge regarding the regulatory mechanisms underlying rapid vacuole degradation and their significance in HECs will be required to further understand HSC emergence. In addition to osmotic cell swelling, vacuoles generally function in autophagic digestion through fusion with lysosomes or direct transition into acidic organelles. Hence, their prospective lytic function should also be assessed.

The biological significance of water transport for cell floating has been suggested in teleost eggs. Intrinsically, high aqueous-content fish eggs are buoyant in sea water, and this increases the chance of oxygen exchange and dispersal over the ocean (Fabra et al., 2005). We found that artificially water-permeated endothelial cells acquire the ability to float in the bloodstream (Fig. 4). Further studies are required to quantitatively determine whether water influx into HECs through AQPs contributes to the buoyancy of HSCs for circulation and subsequent migration into hematopoietic tissues.

In this study, we performed knockout experiments in quail embryos to elucidate the redundant role of AQPs in EHT. CRISPR/Cas9 genome editing in avian embryos by electroporation is effective for knockout analysis in the normal cell population to circumvent lethal effects; however, technically, it does not destroy all genes in the target cell lineages. Therefore, in avian embryos, it was challenging to assess the effect of multiple *AQP* gene knockouts on definitive HSC colony formation in the bone marrow. Further knockout analysis with AQP genes in mouse embryos is needed to elucidate this point.

In summary, this study is the first to demonstrate the involvement of water permeation in EHT using an amniote model system. The findings described here are expected to advance our understanding of embryonic hematopoiesis.

## MATERIALS AND METHODS

### Quail and chicken embryos

Fertilized eggs of tg(pLSi/ΔeGFP) chickens (Motono et al., 2010) were obtained from the Avian Bioscience Research Center at Nagoya University. Tg(tie1:H2B-eYFP) (Sato et al., 2010) and wild-type quail strains were bred in our own quail breeding facility at Kyushu University. Fertilized eggs were incubated at 38°C until electroporation and subsequent analyses. The staging of quail embryos was based on the Hamburger and Hamilton stages of chicken embryos (Hamburger and Hamilton, 1951). The animal study was approved by the Institutional Animal Care and Use Committee of Kyushu University (Authorization number: A20-019).

### Antibodies and immunostaining

An anti-AQP1 polyclonal antibody against the C-terminal 252–270 amino acid sequence, CEEYDLEGDDMNSRVGMKPK, of quail AQP1 (GenBank XM_015852703) was obtained from rabbits using a custom antibody production service (Sigma-Aldrich). For immunostaining, rabbit anti-AQP1 (1:2,000 dilution), QH1 (1:500 dilution, Developmental Studies Hybridoma Bank)(Pardanaud et al., 1987), rabbit anti-Runx1/AML1+Runx3+Runx2 (1:800 dilution, Abcam; we specified immunogen in the dorsal aorta as Runx1), rabbit anti-cleaved Caspase-3 (1:500 dilution, Promega), mouse anti-GFP (1:1,000 dilution, Roche), rabbit anti-GFP (1:1,000 dilution, GeneTex), rabbit anti-mCherry (1:1,000 dilution, GeneTex), rabbit anti-RFP (1:1,000 dilution, Invitrogen), goat anti-RFP (1,000 dilution, Rockland), anti-mouse IgG-AlexaFluor 488 (1:1,000 dilution, Cell Signaling), anti-rabbit IgG-AlexaFluor 488 (1:1,000 dilution, Cell Signaling), anti-mouse IgG-AlexaFluor 594 (1:1,000 dilution, Cell Signaling), anti-rabbit IgG-AlexaFluor 594 (1:1,000 dilution, Cell Signaling), anti-goat IgG-AlexaFluor 594 (1,000 dilution, Abcam), anti-mouse IgG-AlexaFluor 647 (1:1,000 dilution, Cell Signaling), and anti-rabbit IgG-AlexaFluor 647 (1:1,000 dilution, Cell Signaling) were used.

Embryos were dissected at the cervical and flank levels in phosphate-buffered saline (PBS) and fixed in 4% paraformaldehyde (PFA)/PBS overnight at 4°C. The embryos were embedded in 3% agarose/PBS and subjected to vibratome sectioning into 140-μm-thick sections at 5100 mz (Campden Instruments). The slices were pre-blocked with 5% fetal bovine serum (FBS) and 0.2% bovine serum albumin (BSA)/PBST (0.2% Triton X-100 in PBS) for 1 h at 25ºC. They were then incubated overnight at 4ºC with a cocktail of primary antibodies diluted with 5% FBS and 0.2% BSA/PBST. After being washed six times in PBST for 30 min at 25ºC, the slices were incubated overnight at 4ºC with a cocktail of fluorescent secondary antibodies and DRAQ5 (Biostatus) diluted with 5% FBS and 0.2% BSA/PBST. The slices were washed five times in PBST for 30 min at 25ºC, transferred into spacers filled with Fluorokeeper (Nacalai Tesque) on glass slides, mounted with cover glasses, and sealed with nail polish. RapiClear 1.47 (SunJin Lab) was used to clear and mount tg(pLSi/ΔeGFP) chicken-derived specimens. Anti-Runx1 and anti-AQP1 immunostaining were performed sequentially (Fig. 2E).

### Post-embedding immunoelectron microscopy

E3.5 (HH20) quail embryos were fixed in 4% PFA and 0.1% glutaraldehyde/PBS. The embryos were then sectioned in the transverse plane into 200-μm thick sections using a Linear Slicer PRO10 (Dosaka EM). The preparations were dehydrated using increasing concentrations of methanol, embedded in LR Gold resin (Electron Microscopy Sciences), and polymerized under UV lamps at –20°C for 24 h. Ultrathin sections containing the dorsal aorta (80 nm in thickness) were collected on nickel grids coated with a collodion film, rinsed with PBS several times, then incubated with 2% normal goat serum and 2% BSA in 50 mM Tris (hydroxymethyl)aminomethane-buffered saline (TBS; pH 8.2) for 30 min to block non-specific binding. The sections were then incubated with rabbit anti-AQP1 antibody (1:400 dilution) for 1 h at 25ºC. To intensify the detectability of the immunoreaction for AQP1, a streptavidin-biotin intensification kit (Nichirei) was used. Sections were first incubated with biotinylated goat anti-rabbit IgG antibody for 10 min at 25ºC, followed by incubation with avidin-biotin-horse radish peroxidase (HRP) complex solution for 5 min at 25ºC. The sections were then washed with PBS and incubated with goat antibody against HRP conjugated to 6-nm gold particles (1:50 dilution, Jackson ImmunoResearch Laboratory) for 1 h at 25ºC. The sections were contrasted with uranyl acetate and lead citrate and observed using an H-7650 electron microscope (Hitachi) operated at 80 kV.

### cDNA cloning

Total RNA was isolated from E2 (HH12) quail embryos using NucleoSpin RNA Plus (Macherey-Nagel) and reverse-transcribed using PrimeScript II reverse transcriptase (Takara Bio). Full-length cDNA was amplified using Ex Taq (Takara Bio) or PrimesSTAR GXL DNA polymerase (Takara Bio) with the following primers: XhoI-qAQP1 fw: 5’-CGCTCGAGATGGCCAGTGAATTCAAA-3’ and BamHI-qAQP1 rv: 5’-CAGGATCCTTATTTTGGCTTCATTTC-3’; EcoRI-qAQP5 fw: 5’-CCGAATTCATGAAGAGGGAAATATTA-3’ and XbaI-qAQP5 rv: 5’-TATCTAGACTACGGTGGGGTCAACTC-3’; EcoRI-qAQP8 fw: 5’-TCGAATTCGAGATGGAGATGGCAGACTC-3’ and XbaI-qAQP8 rv: 5’-TGTCTAGATACAGCCTCACTTCAGGAAC-3’; XbaI-qAQP9 fw: 5’-TATCTAGAAGCCAGAATGAGCCGGA-3’ and Sal I-qAQP9 rv: 5’-GCGTCGACTTACTACATGTTCGTTAGT-3’. The PCR products were cloned into pBluescript II-SK (pBS, Stratagene). Sequences were determined using the ABI 3130 Genetic Analyzer (Applied Biosystems).

### Plasmid vectors for electroporation

pBS-AQP1-mRFP was generated by inserting the PCR-amplified mRFP derived from pCAGGSY-LifeAct-mRFPruby (Sato et al., 2017) into the 3’ end of AQP1 in pBS-AQP1 (described above) using the In-Fusion Cloning system (Takara Bio) with the following primers: qAQP1-mRFP fw: 5’-aagccaaaagccaccATGGCCAGCTCCGAGGATGT-3’ and mRFP-pBS rv: 5’-gccgctctagaactaTTAAGCGCCTGTGCTATGTC-3’ (the sequences for recombination are indicated in lowercase letters). For the construction of AQP1(R196H)-mRFP, pBS-AQP1-mRFP was linearized at BglII and amplified with PCR primers, including the 196th arginine (R) to histidine (H) mutation: qAQP1 R196H fw: 5’-ccagcacatagttttGGCTCAGCACTGAT-3’ and qAQP1 R196H rv: 5’-aaaactatgtgctggGTTTATTCCACAG-3’. The PCR product was assembled into a circular plasmid, pBS-AQP1(R196H)-mRFP, by In-Fusion Cloning between 15 bp of the 5’ ends (lowercase letters in the primer sequences). pBS-AQP1-2A-eYFP was generated by swapping pBS-AQP1-2A-LifeAct-eYFP and pBS-2A-eYFP at the conserved BamHI and PstI sites. AQP1-mRFP, AQP1(R196H)-mRFP, and AQP1-2A-eYFP in pBS were independently subcloned into pCAGGS (Niwa *et al*., 1991). To construct pCAGGS-H2B-mCherry-2A-AQP1, H2B-mCherry was isolated from pCAGGS-H2B-mCherry by PCR using the following primers (the sequences for recombination are indicated in lowercase letters): pCAG-H2B fw: 5’-atcattttggcaaagtctagATGCCAGAGCCAGCG-3’ and qAQP1-2A rv: gaattcactggccatAGGACCGGGGTTTTCTTCCA-3’, which were inserted into the 5’ end of the AQP1 in pCAGGSY-AQP1 by In-Fusion Cloning. To construct pCAGGS-mRFP, mRFP was isolated from pCAGGSY-LifeAct-mRFPruby (Sato et al., 2017) and subcloned into the pCAGGS vector. For the construction of pCAGGS-mRFP-CAAX, the CAAX sequence was isolated from pCAGGS-mCherry-CAAX by PCR using the following primers (the sequences for recombination are indicated in lowercase letters): mRFP-linker fw: 5’-catagcacaggcgctTCCGGACTCAGATCCACGCG-3’ and pCAGGS-CAAX rv: 5’-tgattcgacgcggccTCAGGAGAGCACACACTTGC-3’, which were inserted into the 3’ end of the mRFP in pCAGGSY-mRFP by In-Fusion Cloning. pCAGGS-eGFP and pCAGGS-Lyn-mCherry have been previously described (Sato et al., 2017). gRNA expression vectors U6.3> gRNA.f+e and U6.3> Control.gRNA.f+e were obtained from Addgene (Gandhi et al., 2017). Oligo DNAs for the gRNA templates were designed as follows (lowercase letters indicate BsaI overhang): qAQP1 exon2 top: 5’-agatGTCTCCATACAGCTTGCAAA-3’ and qAQP1 exon2 bottom: 5’-aaacTTTGCAAGCTGTATGGAGAC-3’; qAQP5 exon1 top: 5’-agatGGCCCACTCCCACCATGAAG-3’ aaacCTTCATGGTGGGAGTGGGCC-3’; and qAQP5 qAQP8 exon1 exon4 bottom: top: 5’-5’-aaacGGGCTAACATGTCTGGACCC-3’ and qAQP8 exon4 bottom: 5’-aaacGGGCTAACATGTCTGGACCC-3’; qAQP9 exon4 top: 5’-agatGAACAGCTGTTGACATCACC-3’ and qAQP9 exon4 bottom: 5’-aaacGGTGATGTCAACAGCTGTTC-3’. Annealed oligos were cloned into U6.3> gRNA.f+e according to Stolfi et al. (Stolfi et al., 2014). pCAGGS-Cas9-2A-mRFP was generated by inserting Cas9-2A derived from pCAG-Cas9-2A-Citrin (obtained from Addgene; Williams et al., 2018) into pCAGGS-mRFP by In-Fusion Cloning with the following primers: MCS-nlsCas9 fw: 5’-tcctcgagtctagatAACGCTAGGCCACCATGGCC-3’ and 2A-linker-mRFP rv: 5’-tcggagctggccatgGATCTGGGCCCGGGA-3’. The plasmids were purified using a NucleoBond Xtra Midi EF kit (Macherey-Nagel) and dissolved in TE (10 mM Tris-HCl pH 8, 1 mM EDTA).

### *In ovo* electroporation

For the overexpression experiments, 5 μg/μL of plasmid solution colored with 2% Fast Green FCF (Nacalai Tesque) was prepared. For the CRISPR/Cas9 genome editing, the plasmid solution was mixed as follows: 0.6 μL of U6.3>qAQP1 exon2 gRNA.f+e (5 μg/μL), 0.6 μL of U6.3>qAQP5 exon1 gRNA.f+e (5 μg/μL), 0.6 μL of U6.3>qAQP8 Exon4 gRNA.f+e (5 μg/μL), 0.6 μL of U6.3>qAQP9 Exon4 gRNA.f+e (5 μg/μL), 1.4 μL of pCAGGS-Cas9-2A-mRFP (5 μg/μL), and 2% Fast Green FCF. To transduce the plasmids into the dorsal aortic roof and floor, somite and lateral plate mesoderm progenitors in the primitive streak at E0.75 (HH4) were targeted. The plasmid solution was injected by a glass capillary and electroporated using a CUY21EX pulse generator (Bex) via the following steps: (i) single poration-pulse at 13 V, 0.08 ms pulse length, and 0.05 ms intervals; (ii) three cycles of driving pulses at 6 V, 50 ms pulse length, and 50 ms intervals.

### *In vitro* EHT

Differentiation of the presomitic mesoderm into HECs was performed *in vitro* according to a previous report by Yvernogeau et al. (Yvernogeau et al., 2016) with slight modifications. Presomitic mesoderm-containing regions were isolated from quail embryos at E2 (HH11-12) using a micro blade K-715 (Feather), followed by brief treatment with Dispase (Wako) in PBS. The presomitic mesoderm was isolated from the surrounding tissues, diced on a silicon-coated Petri dish using tungsten needles, and transferred into collagen-coated glass-bottom dishes (Iwaki) filled with a medium (Yvernogeau et al., 2016). The tissues were cultured for 3 days at 37ºC in a 5% CO2 incubator, and the medium was replaced each day.

### Fluorescence-activated cell sorting (FACS)

Three days after the start of culture, tissues were dispersed in 0.25% trypsin-EDTA/PBS for 2 min at 37°C, resuspended with culture medium, and transferred to 15 mL tubes. After centrifugation and removal of the culture medium, the cells were treated with Cellcover (Anacyte laboratories) for 5 min at 4°C, subsequently incubated with anti-AQP1 polyclonal antibody (1:2000 dilution) and QH1 monoclonal antibody (1:1500 dilution) in Cellcover for 30 min at 4°C. After a brief wash in Cellcover and centrifugation (1,000 rpm, 5 min), the cells were incubated with anti-rabbit IgG Alexa Flour 488 (1:1000 dilution) and anti-mouse IgG Alexa Flour 594 (1:1000 dilution) in Cellcover for 30 min at 4°C. After another brief wash in Cellcover and subsequent centrifugation, the cells were resuspended with Cellcover to a 1 × 10^7^ cells/mL concentration. AQP1+/QH1+ and AQP1-/QH1- cells were sorted by FACS Aria SORP (Becton Dickinson) and collected into Trizol LS (Ambion). Total RNA was purified according to the Trizol LS protocol.

### RNA-seq analysis

After amplification of the cDNA library using the Smart-Seq v4 Ultra Low Input Kit (Clontech), the library was prepared using the NEBNext Ultra RNA library prep kit for Illumina (New England Biolabs). Sequences with 150 bp read length × 2 paired ends and 6 Gb of data were obtained using the next-generation sequencer, NovaSeq 6000 (Illumina). Sequence products were quality-checked using the software FastQC (Cock et al., 2009), and low-quality sequence ends were removed using Trimmomatic (Bolger *et al*., 2014). The sequence was then mapped to the Japanese quail reference genome (*Coturnix japonica*, https://www.ncbi.nlm.nih.gov/assembly/gcf_001577835.2) using HISAT2 (Kim *et al*., 2015). The number of raw reads mapped to known exon regions was calculated using featureCount (Liao *et al*., 2014), and then TMP (transcripts per million) values were calculated. The software edgeR (Robinson *et al*., 2009) was used to calculate the expression variation between samples after normalization using the TMM (trimmed mean of M value) method based on read count values. For gene groups that met the conditions of expression difference log2 FC (fold change) > 1 and P value < 0.05, MA plots were performed using R to visualize the gene groups with up- and down-regulated expression, respectively. Gene ontology (GO) analyses were performed by Metascape (Zhou et al., 2019) using GO terms for *H. sapiens*.

### Analysis of CRISPR/Cas9 genome editing

Regions with increased expression of mRFP from the three embryos were collected 28 h after electroporation and lysed in 0.15 M NaCl, 10 mM Tris-HCl (pH 8), 10 mM EDTA, and 0.1% sodium dodecyl sulfate (SDS). Genome fragments of each AQP gene were PCR-amplified using MightyAmp DNA polymerase ver.3 (Takara Bio) with the following primers: qAQP1 intron1 fw: 5’-ATGATGGTAGAGGCCTGAAC-3’ and qAQP1 618 rv: 5’-GTTGTTGGCAATCAGTGCTG-3’; qAQP5 exon1 −117 fw: 5’-GCTGCTGTTGCATATATAGA-3’ and qAQP5 exon1 +272 rv: 5’-CACATAGAAGAGTGTCCGGA-3’; qAQP8 intron3 fw: 5’-TCACCAGCCTCTGCAATTAC-3’ and qAQP8 intron4 rv: 5’-TATGAAGCCAGCAGACATGC-3’; qAQP9 intron 3 fw: 5’-TGTGAACTGGAGAAGCACTG-3’ and qAQP9 intron5 rv: 5’-CAGTACCTAGAGCATGTCTG-3’. The PCR products were cloned into pCR4-TOPO (Invitrogen) or pBS for sequencing. Sequence alignment was performed using Genetyx software (Genetyx).

### Time-lapse microscopy

Electroporated quail embryos were developed *in ovo* until E2 (HH12) and transferred to *ex ovo* as previously described (Sato and Lansford, 2013). Time-lapse imaging of the quail embryos (Figure 3B and C, Movies 1 and 2) was performed using an upright confocal laser microscope, Fluoview 1200 (Olympus), equipped with a GaAsP detector and thermal stage (Tokai Hit). The embryos were placed in glass-bottom dishes (Iwaki) coated with agar-albumen media (Chapman et al., 2001) and placed with the glass-bottom side up on the microscopic stage to set the embryos within working distance. Fluorescent signals were imaged using a UPLASPO 30× N.A. 1.05 silicon immersion objective lens with a 235-μm pinhole size, 8-μs/pixel scan speed, 640 × 640 resolution, 1.80-μm z-intervals, and 6-min time intervals. The drifting of the images was corrected using Imaris (Bitplane) as described by Sato and Lansford (Sato and Lansford, 2013). For imaging at E3 (Movies 3), electroporated quail embryos were developed *in ovo* until E2 (HH12), transferred to plastic hemispherical cups (40 mm in diameter) with the whole-egg content, and placed inside custom glass-top plastic chambers humidified with sterilized water. Video rate imaging were performed using an epifluorescence stereomicroscope MVX10 (Olympus) equipped with a custom outer thermal chamber and heating unit SSH300 (Shoei Shokai). Fluorescent signals were imaged using MVPLAPO 2xC lens, 3.2× zoom, 512 × 512 resolution, single plane, and stream acquisition by ORCA-Flash 4.0.

### Imaging of immunostained specimens

Immunostained embryo slices and cultured tissues were imaged using Fluoview 1200 through UPLSAPO 30× N.A. 1.05 silicon-, UPLSAPO 60× N.A. 1.30 silicon-, and UPLXAPO 100× N.A. 1.47 oil-immersion objective lenses at optimal configurations. For the quantification of apoptotic cells (Figure S3A and B), whole-mount embryos, immunostained with anti-cleaved Caspase-3 antibody, were imaged using UPLSAPO 10x N.A. 0.40 objective lens at optimal configurations.

### *In situ* hybridization

Digoxigenin (DIG)-labeled antisense RNA probes for AQP1, AQP5, AQP8, and AQP9 were prepared according to the manufacturer’s instructions (Roche). *In situ* hybridization was performed as follows: E3.5 (HH20) quail embryos were fixed in 4% PFA/PBS overnight at 4ºC and embedded in paraffin wax. The sections (12-μm thick) were de-paraffinized twice in xylene for 5 min and rehydrated in a graded ethanol series (twice at 100%, then 90% and 70%; 5 min for each step). After being washed twice with PBST (0.1% Tween 20/PBS) for 5 min, the sections were treated with 2 μg/mL proteinase K/PBST for 7 min at 37ºC, followed by refixation in 4% PFA/PBS for 20 min at 25ºC. The sections were washed thrice in PBST for 5 min and incubated with 50% formamide, 5× SSC (0.6 M NaCl, 0.06 M tri-sodium citrate dehydrate, pH 5), and 0.5% SDS for 10 min at 25ºC. They were then hybridized with the DIG-labeled probes in ULTRAhyb (Invitrogen) overnight at 65ºC. The sections were washed in wash solution 1 (50% formamide, 5× SSC, and 1% SDS) for 30 min at 65°C, twice in wash solution 2 (50% formamide and 2× SSC) for 30 min each at 65°C, and once in a 1:1 mixture of wash solution 2 and TBST (0.8% NaCl, 0.02% KCl, 25 mM Tris-HCl pH 7.5, and 0.1% Tween 20) for 5 min at 65°C. After washing thrice in TBST for 5 min at 25ºC, the sections were pre-blocked with 2% blocking reagent (Roche)/20% FBS/TBST at 25ºC for 1 h, followed by incubation with anti-DIG-alkaline phosphatase-Fab fragments (1:1000 dilution, Roche) overnight at 4 °C. The sections were washed four times for 5 min in 2 mM levamisole/TBST and for 5 min in NTMT (50 mM MgCl2, 100 mM NaCl, 100 mM Tris-HCl pH 9.5, and 0.1% Tween 20). They were then incubated with 100 μg/mL NBT (Roche) and 37.5 μg/mL BCIP (Roche) in NTMT at 25ºC for 6–8 h. After the color reaction, the sections were washed in water for 5 min and dehydrated in a graded ethanol series (50%, 70%, and 90%, 5 min in each step, and twice at 100% for 5 min). The sections were immersed in xylene twice for 5 min and sealed with cover glasses using mounting media. Images were captured using a BZ-X700 microscope (Keyence).

### Image segmentation and quantification

The fluorescent signals of Runx1 and DRAQ5 in a constant xyz field size (150.45 × 150.45 × 40.00 μm^3^) of the tg(pLSi/ΔeGFP) chick embryo slices were automatically segmented and counted by Imaris (Fig. 1E). Vacuolated cells were manually marked on the Imaris, and their numbers were counted (Fig. 1F). To measure cell roundness, optically sectioned eGFP and mRFP images were converted to 8-bit images, threshold levels were adjusted, and particle analyses were performed using ImageJ (NIH). Inverted images were used to measure the vacuole area sizes (Fig. 1G and H). Fluorescently inseparable cell clusters and cavities were eliminated from the measurement candidates. In cases where there were multiple vacuoles inside a single cell section, the area sizes were totaled. Runx1-positive HECs were selected for measurement (Fig. 6D–F). Nuclear eYFP signals in tg(tie1:H2B-eYFP) embryos (Fig. S6) were used for the automatic detection and counting of endothelial cells by Imaris (Fig. 6C and G). Cleaved caspase-3 signals in a constant xyz field size, 1271.2 × 1271.2 × 80.5 μm^3^, were automatically detected and counted by Imaris (Fig. S2A and B).

### Statistical analyses

The number of analyzed embryos (n) is stated in the figure legends. Graph drawing and statistical analyses were performed using GraphPad Prism (GraphPad software). Unpaired *t*-tests and one-way ANOVA were employed for the comparison of two and multiple groups, respectively. Bars in Figs 1E and F, 3C and D, 6D–H, and S2B, and horizontal lines in Fig. 1H graphs indicate the mean values.

## Acknowledgments

We thank the Avian Bioscience Research Center at Nagoya University, the Research Center for Human Disease Modeling at Kyushu University, David Huss and Rusty Lansford (Children’s Hospital Los Angeles, University of Southern California), Kenji Miyagawa, Xinyi Wang, and Takashi Miura (Kyushu University), and Tomohiro Kawahara (Kyushu Institute of Technology) for their help and guidance.

## Author contributions

Y.S. designed and performed the experiments, analyzed the data, and wrote the manuscript. M.S. and C.T. performed preliminary analyses. M.S.K. performed FACS and RNA-seq. S.M. and H.S. performed immunoelectron microscopy.

## Data availability

Datasets of RNA-seq will be deposited to the Gene Expression Omnibus (GEO) repository on publication.

